# Environmental enrichment selectively restores brain metabolic activity during cocaine abstinence

**DOI:** 10.1101/2025.04.26.650753

**Authors:** Pauline Belujon, Virginie Lardeux, Emilie Dugast, Sophie Sérrière, Sylvie Bodard, Julie Busson, Clovis Tauber, Laurent Galineau, Marcello Solinas

## Abstract

**Background:** Environmental enrichment (EE) is a promising strategy to promote recovery from addiction, but its neurobiological mechanisms remain poorly understood. This study investigates how exposure to EE during abstinence dynamically affects brain neuroadaptations induced by voluntary intake of cocaine.

**Methods:** Using longitudinal 18FDG microPET imaging, we examined brain metabolic activity in rats following extended access cocaine self-administration. After establishing escalation of cocaine intake, rats were housed in either enriched or standard environments during a four-week abstinence period. Brain metabolic activity was assessed after one and four weeks of abstinence.

**Results:** Cocaine self-administration produced widespread decreases in cortical metabolic activity, particularly in regions involved in executive function (orbitofrontal cortex, anterior cingulate) and interoception (insula), while increasing activity in emotional and motivational circuits (nucleus accumbens, amygdala, mesencephalon). EE selectively normalized these alterations through temporally distinct mechanisms: rapidly restoring nucleus accumbens and amygdala function while gradually affecting prefrontal cortical activity. After four weeks, rats housed in enriched environments showed significantly normalized metabolic activity in the orbitofrontal cortex and dorsolateral striatum compared to those in standard housing, with persistent changes in anterior cingulate cortex and ventral posterior hippocampus.

**Conclusions:** Our findings reveal circuit-specific and temporally distinct effects of environmental enrichment on cocaine-induced brain alterations. These insights could inform the development of more targeted therapeutic approaches for addiction recovery.

## Introduction

One of the hallmarks of addiction and one of its most troubling aspects is the long-lasting risk of relapse even after protracted periods of abstinence (1–3). Over the past decades, neuroimaging studies in humans and mechanistic studies in animals have demonstrated that drugs of abuse produce persistent plastic changes in the brain that are thought to underlie the chronic nature of addiction (4,5). Based on clinical and brain-imaging data in humans, Goldstein and Volkow proposed the Impaired Response Inhibition and Salience Attribution (I-RISA) model that postulates that addiction results from dysregulation in the coordinated activity of different brain circuits (6,7). In this model, reduced activity in prefrontal and frontal cortical areas leads to deficits in executive function and response inhibition, while increased reactivity in limbic regions leads to heightened responsivity to drug-related cues and stress (6,7). In particular, studies in humans with cocaine addiction have shown hypoactivity in the anterior cingulate and orbitofrontal cortex associated with decision-making deficits (8,9), and hyperactivity in regions like the amygdala associated with increased stress reactivity (10).

Animal studies have been essential in understanding the mechanisms underlying these brain changes (4,11,12). In the last decades, brain imaging studies have provided important insights into the changes induced by acute and chronic intake of drugs (13–20) and the neuroadaptations that persist after discontinuation of drug use which could be associated with incubation of craving and long term risks of relapse (21–26). In particular, using microPET imaging in rats, we have shown that extended access to cocaine self-administration produces persistent changes in brain metabolic activity that strikingly parallel those described in humans (24). Cocaine-exposed rats show decreased activity in cortical regions involved in executive function and decision-making, and increased activity in regions involved in motivation, memory, stress and emotional processing (24). Despite our growing understanding of cocaine-induced brain changes, developing effective treatments for addiction remains challenging.

Among the novel strategies to help recovering from addiction, environmental enrichment has emerged as a promising therapeutic approach (27–30). Housing animals in stimulating environments that provide opportunities for social interaction, physical activity, and cognitive engagement during periods of drug abstinence reduces drug seeking and taking behaviors in animal models of addiction (29,31). While a few studies have investigated neurobiological mechanisms associated with the beneficial effects of EE on relapse (30,32–34), the mechanisms by which environmental enrichment produces its therapeutic effects on the brain and their dynamics remain poorly understood.

In this study, we used longitudinal 18FDG microPET imaging to investigate how exposure to environmental enrichment during abstinence affects cocaine-induced changes in basal brain metabolic activity. We first allowed rats to self-administer cocaine under extended access conditions that lead to escalation of intake (35) and then housed them in either enriched or standard environments during abstinence. By comparing brain activity patterns across groups and time points, we aimed to better understand how environmental enrichment influences the brain changes associated with cocaine addiction and withdrawal.

## Methods

### Subjects and experimental design

Adult (11-12 weeks of age) male Sprague-Dawley rats (Janvier, France), experimentally naive at the start of the study, were housed in a temperature- and humidity-controlled room and maintained on a 12-h light/dark cycle (light on at 7.00 AM). All experiments were conducted in accordance with European Union directives (2010/63/EU) for the care of laboratory animals approved by the local ethics committee (COMETHEA, CEEA Val de Loire, #2015033014555999).

### Housing Conditions

On arrival, rats were housed two per cage for about 1 week before intrajugular catheterization surgery. Rats were anaesthetized using isoflurane (5% induction, 2.5% surgery) and implanted with catheters into the right jugular vein. Animals were allowed to recover for 5-7 days before the self-administration sessions began. After surgery, rats were housed individually during the entire period of active self-administration. At the end of the last self-administration session, rats were randomly divided into two groups, one housed in SE and the other in EE conditions assuring that similar levels of drug self-administration were observed in the two groups. SE and EE conditions were the same as previously used (36–38). For SE, rats were housed in groups of three in cages sized 60 × 38 × 20□cm. For EE, rats were housed in groups of three in cages sized 80 × 50 × 100□cm. Each EE cage contained a house, a running wheel, three floors connected by ramps or tunnels and four toys that were changed once per week. Rats were housed in the animal facility in SE or EE housing conditions during a 4-week period of abstinence. A total of four experimental groups were obtained: NaiveSE, NaïveEE, CocSE and CocEE. The timeline of the experiment, including the different experimental groups, is illustrated in Figure 1A.

**Figure 1.**
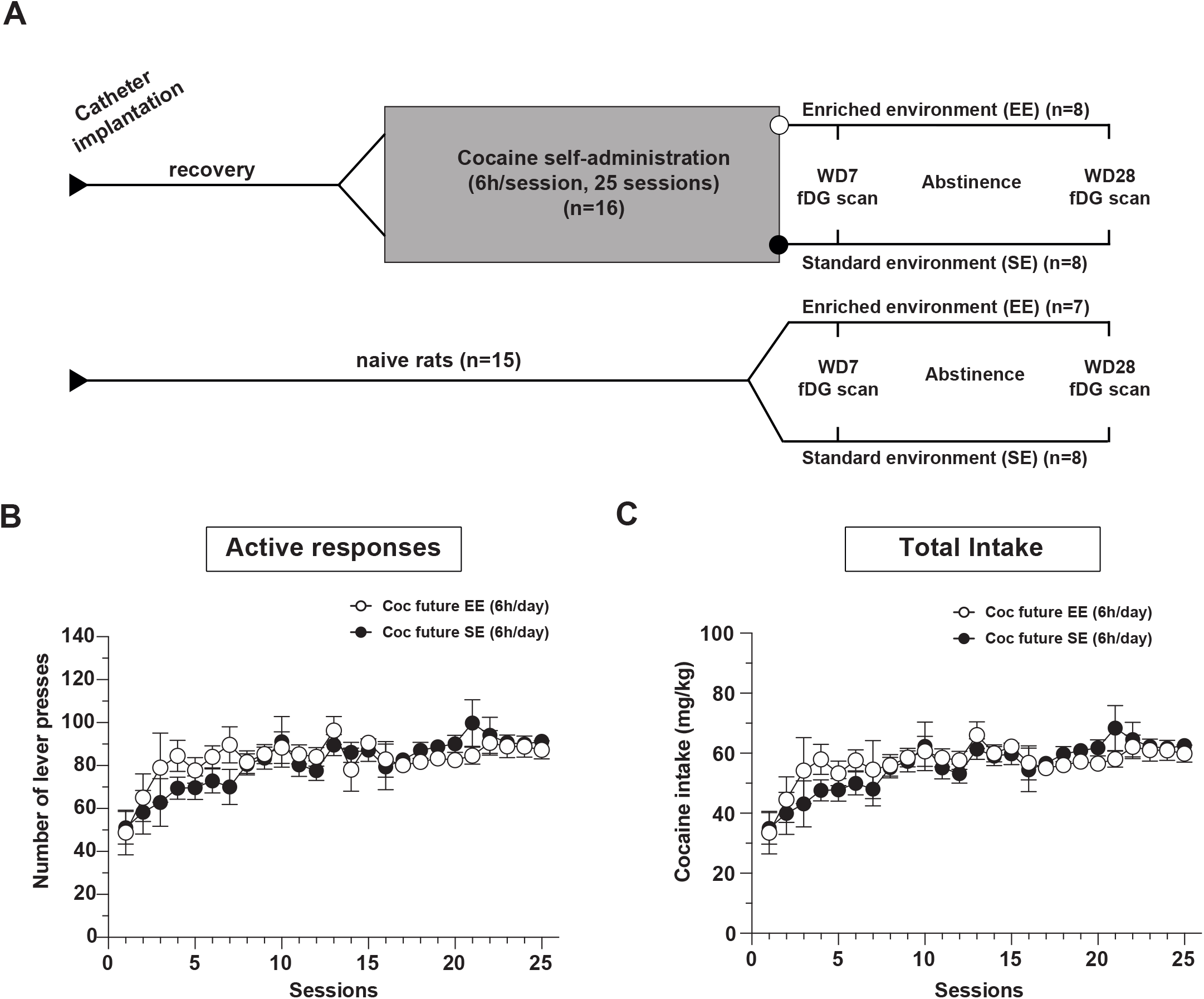
General experimental design and levels of self-administration in rats that were housed in SE or EE during abstinence. A) Cocaine rats were allowed to self-administer cocaine for 25 × 6h sessions and then pseudo-randomly assigned to SE or EE conditions assuring similar levels of cocaine intake. 18FDG scan were performed after 1 week and 4 weeks of abstinence. Naive control animals were age-matched rats that were house similarly to cocaine rats but did not undergo surgery and self-administration procedures. B) Number of cocaine injections/sessions in future SE and future EE animals. C) Cocaine Intake in future SE and future EE animals. Notice that cocaine intake did not differ between the two groups.

### Cocaine self-administration apparatus and procedure

#### Catheter implantation

Rats were prepared for cocaine self-administration by surgical catheterization of the right jugular vein. Briefly, rats were anesthetized with isoflurane (5% induction, 2.5% maintenance) in O2 and administered with the nonsteroidal anti-inflammatory ketoprofen (2.5mg/kg, s.c.). Animals were placed on a heating pad from the induction of the anesthesia to the end of the surgery. A handmade silastic catheter was inserted into the jugular vein and the distal end was led to the back between the scapulae. Rats were allowed to recover for 7 days and flushed daily with 0.1ml sterile saline (0.9%), gentamicin (20mg/ml) and heparin (100 UI/mL) in sterile saline to help protect against infection and catheter occlusion.

#### Apparatus

Experiments were conducted in MedAssociates operant-conditioning chambers, equipped with retractable levers as operanda, a cue-light above the active lever, a house light and controlled by MedAssociates interfaces and MED-PC IV software (www.medassociates.com).

#### Cocaine self-administration procedure

Rats were allowed to self-administer cocaine (Cooper, France; 6g/L in saline solution 0.9%; 0.75mg/kg/infusion) for 6h/day for 25 sessions, using a Fixed Ratio 1 (FR1) schedule of reinforcement. A single press on the active lever resulted in one intravenous (i.v.) cocaine infusion with the concomitant activation of the light that remained on for 5s and then pulsed for 5s, followed by a 5s time-out. Inactive lever presses were recorded but did not produce any consequences.

At the end of the last self-administration session, rats were transferred from the animal facility at the University of Poitiers to the animal facility at University of Tours by an authorized transporter, placed in the SE or EE previously described and underwent abstinence for 4 weeks (EE: N=8 and SE: N=8). Metabolic imaging using 18FDG was performed in the same rats after 6–8 days (1 week) and 27–29 days (4 weeks) of abstinence. Naive rats of the same age and with analogous housing conditions but that did not undergo self-administration were used as controls (EE: N=7 and SE: N=8).

### Brain Imaging

Local uptake of 18FDG reflects cerebral metabolic rates of glucose utilization and allows the investigation of regional brain metabolic status (39). Metabolic imaging using 18FDG was performed under basal conditions using the same procedure as previously described (24). Briefly, rats were habituated to the PET experimental procedures for 4 days before each scan and fasted overnight before each scan. The day of brain-imaging acquisition, awake rats were injected with 18FDG (18.5 MBq/100 g i.p.; Cyclopharma, Tours), and placed in the habituation cage for 45 min. Then, they were anesthetized using isoflurane 4% (Baxter, Maurepas, France), placed on a heating pad (Minerve, Esternay, France) and centered in the field of view of the Explore VISTA-CT microPET camera (GE Healthcare, Velizy, France). CT-scan was performed for attenuation correction of PET images and a list-mode PET acquisition of 30 min started 60 min after 18FDG injection. After data reconstruction using a 2-D OSEM algorithm, all images were co-registered and normalized for tissue activity in the whole brain. Quantitative results were expressed as mean ± SD and were presented on Z-score maps (for more details, refer to (40)). Analyses focused on brain areas known to be key nodes in addiction and that exhibited changes in 18FDG uptake during cocaine abstinence: the cingulate (Cg), orbitofrontal (OFC), prelimbic/infralimbic (PrL/IL), insular and motor cortices, in addition to the dorsal striatum (DStr), nucleus accumbens (NAc), substantia nigra/ventral tegmental area (SN/VTA), amygdala (Amyg), and hippocampus (Hipp) (24,41).

### Statistical analysis

For self-administration, data were analyzed by two-way repeated measures ANOVA with time designated as a within-subject factor and future environment exposure (Coc SE or Coc EE) as a between-subject factor.

For micro-PET data, a voxel-based analysis was used to assess the differences in cerebral 18FDG uptake between the averaged brains of cocaine vs control rats at each stage of abstinence for rats housed in EE vs SE. The regions of interest were derived from Schiffer’s templates (42) using PMOD v3.2 software (PMOD Technologies Ltd, Switzerland) and applied to Z-score and effect size maps to obtain the Z-score and *d* values in these areas (40,43). Statistical analyses focused on inter-group comparisons investigating differences in metabolic activity between (i) cocaine and naïve rats housed in each environment at each time point, and (ii) EE vs SE in cocaine and naïve rats at each time point using a two-tail unpaired Student t-test. Differences were considered significant when p<0.05 for signals of at least 50 contiguous voxels, with *d* values of at least 0.80.

## Results

### Cocaine self-administration

Prior to being assigned to different housing conditions, all rats showed similar patterns of cocaine self-administration (Figure 1B, C). Statistical analysis revealed no significant difference between future SE and future EE rats in the number of active lever presses (two-way ANOVA: time, F_(24,360)_=5.81, p<0.05 ; group SE/EE, F_(1,14)_=0.1150, p=0.7395 ; interaction, F_(24,336)_=1.006, p=0.4582) or in cocaine intake (two-way ANOVA: time, F_(24,336)_=5.777,p<0.05 ; group SE/EE, F_(1,14)_=0.1147, p=0.7399 ; interaction, F_(24,336)_=0.7573, p=0.7896).

### Effects of Environmental Enrichment on Brain Activity

First, we investigated the effects of EE in naive rats (NaiveSE vs NaiveEE). One week of EE produced specific changes in brain metabolic activity (Table 1 and Figure 2). Decreased activity was observed in the anterior cingulate cortex (ACC) (p=0.0149), motor cortex (p=0.0018), insula (p=0.0019), anterior hippocampus (p=0.0104), and SN/VTA (p=0.0002). Increased activity was found in the PrL/IL (p=0.0047), dorsomedial and dorsolateral striatum (p=0.0011 and p=0.0021), and posterior Hipp (dorsal: p=0.0043; ventral: p=0.0067). After four weeks, most changes normalized except for persistent alterations in motor cortex (increased, p=0.0081), insula (decreased, p=0.0220), ventral posterior hippocampus (increased, p=0.0011), and SN/VTA (decreased, p=0.0085).

**Table 1:** Statistically significant differences in 18FDG uptake between naïve rats housed for one and four weeks in EE vs SE (NaiveEE vs NaiveSE).

|  | 1 week |  |  | 4 weeks |  |  |
| --- | --- | --- | --- | --- | --- | --- |
|  | Z-score | d-value | p-value | Z-score | d-value | p-value |
| ACC | -2.64±0.35 | 1.14 | p=0.0149 | — | — | — |
| Mot | -2.62±0.55 | 1.13 | p=0.0018 | 2.47±0.50 | 1.09 | p=0.0081 |
| OFC | — | — | — | — | — | — |
| PrL/IL | 2.73±0.40 | 1.16 | p=0.0047 | — | — | — |
| Insula | -3.10±0.81 | 1.24 | p=0.0019 | -2.55±0.48 | 1.10 | p=0.0220 |
| DMStr | 2.88±0.71 | 1.21 | p=0.0011 | — | — | — |
| DLStr | 2.78±0.66 | 1.19 | p=0.0021 | — | — | — |
| NAc | — | — | — | — | — | — |
| Amyg | — | — | — | — | — | — |
| Ant. Hipp | -2.49±0.52 | 1.11 | p=0.0104 | — | — | — |
| Dors. Post. Hipp | 2.69±0.61 | 1.15 | p=0.0043 | — | — | — |
| Vent. Post. Hipp | 2.56±0.41 | 1.11 | p=0.0067 | 2.81±0.54 | 1.16 | p=0.0011 |
| SN/VTA | -3.11±1.05 | 1.22 | p=0.0002 | -2.52±0.25 | 1.09 | p=0.0085 |

ACC, anterior cingulate cortex; Amyg, amygdala; Ant. Hipp, anterior hippocampus; DLStr, dorsolateral striatum; DMStr, dorsomedial striatum; Dors. Post. Hipp, dorsal posterior hippocampus; Mot, motor cortex; NAc, nucleus accumbens; OFC, orbitofrontal cortex; PrL/IL, prelimbic/infralimbic cortices; SN/VTA, substantia nigra/ventral tegmental area; Vent. Post. Hipp, ventral posterior hippocampus.

**Figure 2.**
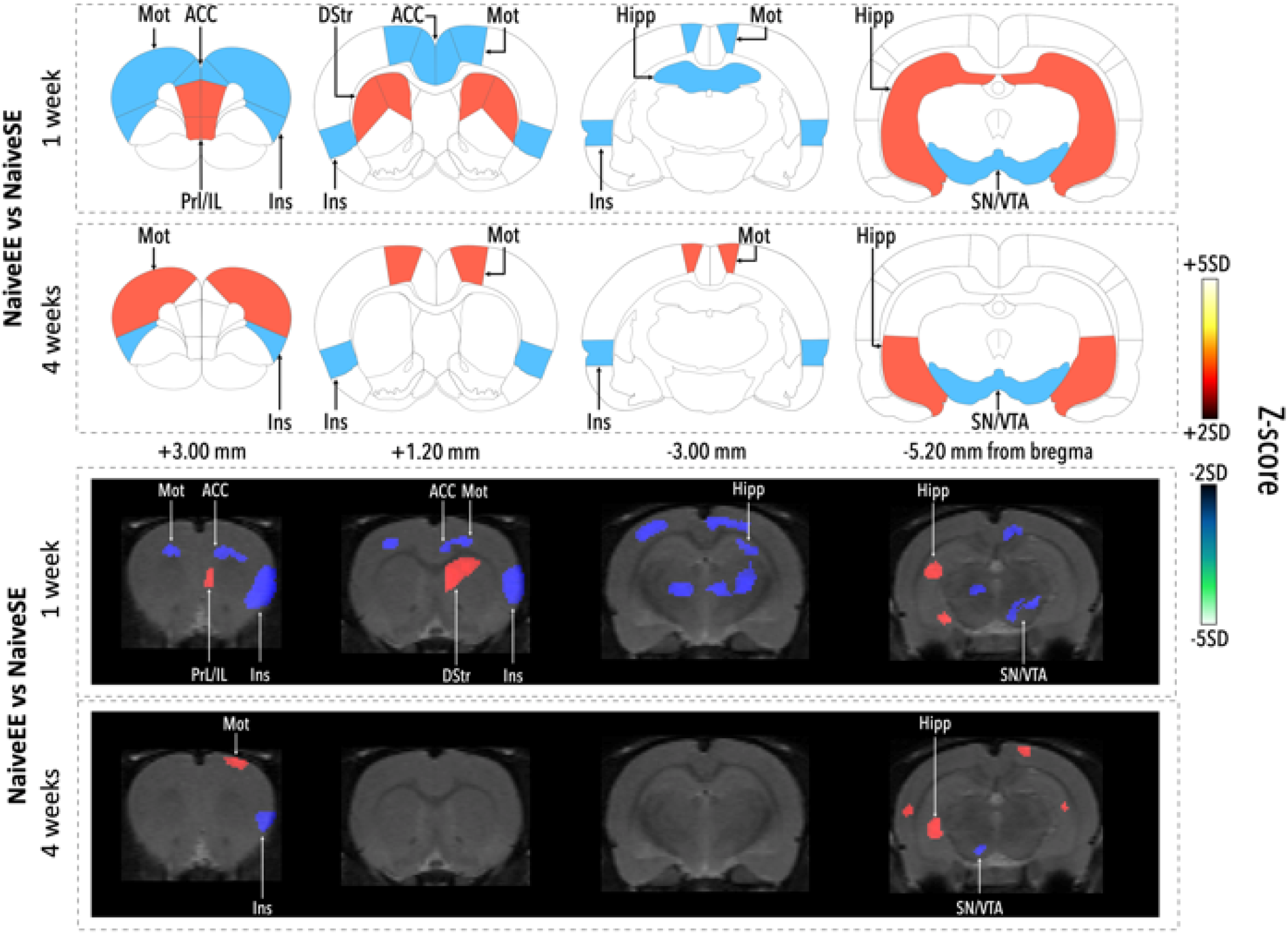
Effects of one and four weeks of exposure to EE on brain metabolic activity. Changes in metabolic activity in naive rats housed in environmental enrichment (NaiveEE) compared to naïve rats housed in standard environment (NaïveSE) presented on representative coronal plates of the Paxinos and Watson atlas (upper panel), and on coronal images of z-score maps fused with an MRI template (lower panel). Increases in 18FDG uptake from dark red to yellow, decreases in 18FDG uptake from black to light blue; Student’s two-tailed t-test. ACC, anterior cingulate cortex; DStr, dorsal striatum; Hipp, hippocampus; Ins, insula; Mot, motor cortex; PrL/IL, prelimbic/infralimbic cortices; SN/VTA: substantia nigra/ventral tegmental area.

### Effects of Cocaine on Brain Metabolic Activity During Abstinence

Then, we verified that we could replicate our findings of cocaine-induced changed in metabolic activity (24). Consistent with our previous findings, in rats housed in standard environments (CocSE vs NaiveSE), one week of abstinence was characterized by widespread decreases in metabolic activity (Table 2 and Figure 3), particularly in cortical regions including the ACC (p=0.0018), motor cortex (p<0.0001), OFC (p<0.0001), PrL/IL (p=0.0011), and insula (p=0.0017). The Nac also showed decreased activity (p=0.002). In contrast, increased activity was observed in the dorsomedial striatum (p=0.0242), posterior hippocampus (dorsal: p=0.0155; ventral: p=0.0038), and SN/VTA (p=0.0396). After four weeks of abstinence, while some brain regions showed recovery (PrL/IL, NAc, and the motor cortex), persistent decreases were observed in the ACC (p=0.0362), OFC (p=0.0002), insula (p=0.0018), and dorsolateral striatum (p=0.012). The ventral posterior hippocampus (p=0.0008) and SN/VTA (p=0.0325) maintained increased activity. Finally, the anterior hippocampus show switch from decrease to increase in metabolic activity (p=0.0074).

**Table 2.** Statistically significant differences in 18FDG uptake between cocaine self-administered and naïve rats housed in SE for one and four weeks of abstinence (CocSE vs NaiveSE).

|  | 1 week |  |  | 4 weeks |  |  |
| --- | --- | --- | --- | --- | --- | --- |
|  | *Z*-score | *d*-value | *p*-value | *Z*-score | *d*-value | *p*-value |
| ACC | -2.91±0.71 | 1.14 | p=0.0018 | -2.30±0.12 | 1.00 | p=0.0362 |
| Mot | -3.22±0.86 | 1.22 | P<0.0001 | — | — | — |
| OFC | -3.49±0.95 | 1.26 | P<0.0001 | -3.16±0.79 | 1.21 | p=0.0002 |
| PrL/IL | -2.77±0.52 | 1.12 | p=0.0011 | — | — | — |
| Insula | -2.91±0.83 | 1.17 | p=0.0017 | -2.98±0.89 | 1.17 | p=0.0018 |
| DMStr | 2.33±0.14 | 1.00 | p=0.0242 | — | — | — |
| DLStr | — | — | — | -2.82±0.75 | 1.15 | p=0.012 |
| NAc | -3.17±0.74 | 1.22 | p=0.002 | — | — | — |
| Amyg | — | — | — | — | — | — |
| Ant. Hipp | -2.55±0.25 | 1.07 | p=0.0080 | 2.25±0.07 | 0.98 | p=0.0074 |
| Dors. Post. Hipp | 2.41±0.61 | 1.05 | p=0.0155 | — | — | — |
| Vent. Post. Hipp | 2.46±0.39 | 1.05 | p=0.0038 | 2.84±0.51 | 1.14 | p=0.0008 |
| SN/VTA | 2.48±0.36 | 1.04 | p=0.0396 | 2.35±0.18 | 1.00 | p=0.0325 |

ACC, anterior cingulate cortex; Amyg, amygdala; Ant. Hipp, anterior hippocampus; DLStr, dorsolateral striatum; DMStr, dorsomedial striatum; Dors. Post. Hipp, dorsal posterior hippocampus; Mot, motor cortex; NAc, nucleus accumbens; OFC, orbitofrontal cortex; PrL/IL, prelimbic/infralimbic cortices; SN/VTA, substantia nigra/ventral tegmental area; Vent. Post. Hipp, ventral posterior hippocampus.

**Figure 3.**
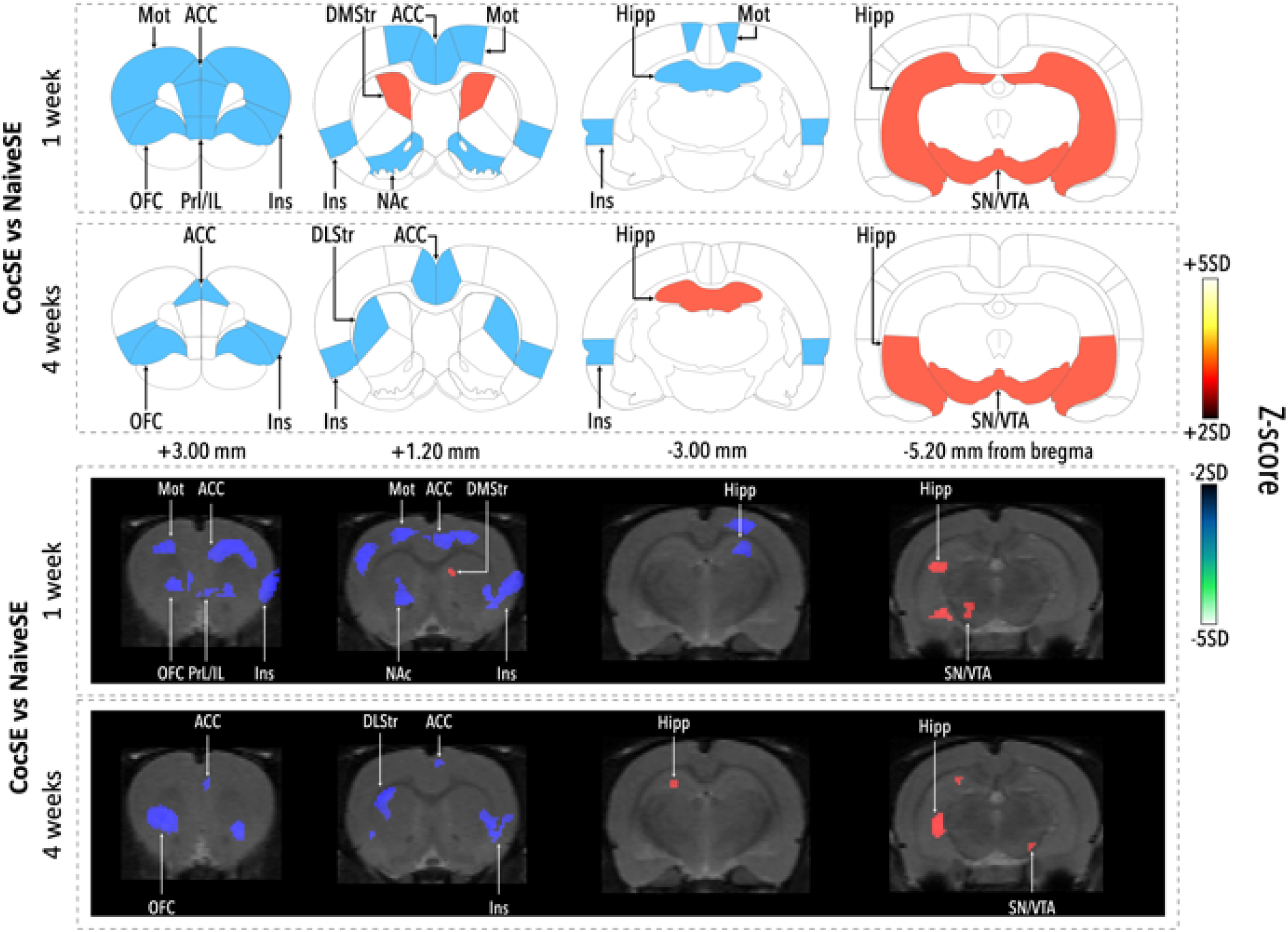
Effects of cocaine self-administration on brain metabolic activity after one and four weeks of housing in SE. Changes in metabolic activity in rats that self-administered cocaine (CocSE) compared to naïve rats (NaïveSE) housed in standard environments during abstinence presented on representative coronal plates of the Paxinos and Watson atlas (upper panel), and on coronal images of z-score maps fused with an MRI template (lower panel). Increases in 18FDG uptake from dark red to yellow, decreases in 18FDG uptake from black to light blue; Student’s two-tailed t-test; p < 0.01. ACC, anterior cingulate cortex; DLStr, dorsolateral striatum; DMStr, dorsomedial striatum; Hipp, hippocampus; Ins, insula; Mot, motor cortex; NAc, nucleus accumbens; OFC, orbitofrontal cortex; PrL/IL, prelimbic/infralimbic cortices; SN/VTA: substantia nigra/ventral tegmental area.

### Effects of Environmental Enrichment on Cocaine-Induced Brain Changes

When cocaine-exposed rats were housed in enriched environments (CocEE vs NaiveEE), changes in metabolic were found only in a few regions (Table 3 and Figure 4). Indeed, after one week, changes were limited to decreases in the ACC (p=0.0160), PrL/IL (p=0.0104), amygdala (p=0.0035), and anterior hippocampus (p=0.0071), with increases in NAc (p=0.0006) and posterior hippocampus (dorsal: P=0.0032; ventral: p=0.0138). After four weeks, we found significant increases in ventral posterior hippocampus (p=0.0176) and significant decreases in the ACC (p=0.0049) and in the motor cortex (p=0.0004).

**Table 3.** Statistically significant differences in 18FDG uptake between cocaine self-administered and naïve rats housed in EE for one and four weeks of abstinence (CocEE vs NaiveEE).

|  | 1 week |  |  | 4 weeks |  |  |
| --- | --- | --- | --- | --- | --- | --- |
|  | Z-score | d-value | p-value | Z-score | d-value | p-value |
| ACC | -2.38±0.16 | 1.04 | p=0.0160 | -2.68±0.35 | 1.15 | P=0.0049 |
| Mot | — | — | — | -2.81±0.96 | 1.19 | p=0.0004 |
| OFC | — | — | — | — | — | — |
| PrL/IL | -2.63±0.42 | 1.12 | p=0.0104 | — | — | — |
| Insula | — | — | — | — | — | — |
| DMStr | — | — | — | — | — | — |
| DLStr | — | — | — | — | — | — |
| NAc | 2.64±0.46 | 1.12 | p=0.0006 | — | — | — |
| Amyg | -2.84±0.73 | 1.16 | p=0.0035 | — | — | — |
| Ant. Hipp | -2.70±0.45 | 1.13 | p=0.0071 | — | — | — |
| Dors. Post. Hipp | 2.60±0.58 | 1.12 | P=0.0032 | — | — | — |
| Vent. Post. Hipp | 2.66±0.33 | 1.13 | p=0.0138 | 2.51±0.36 | 1.11 | p=0.0176 |
| SN/VTA | — | — | — | — | — | — |

ACC, anterior cingulate cortex; Amyg, amygdala; Ant. Hipp, anterior hippocampus; DLStr, dorsolateral striatum; DMStr, dorsomedial striatum; Dors. Post. Hipp, dorsal posterior hippocampus; Mot, motor cortex; NAc, nucleus accumbens; OFC, orbitofrontal cortex; PrL/IL, prelimbic/infralimbic cortices; SN/VTA, substantia nigra/ventral tegmental area; Vent. Post. Hipp, ventral posterior hippocampus.

**Figure 4.**
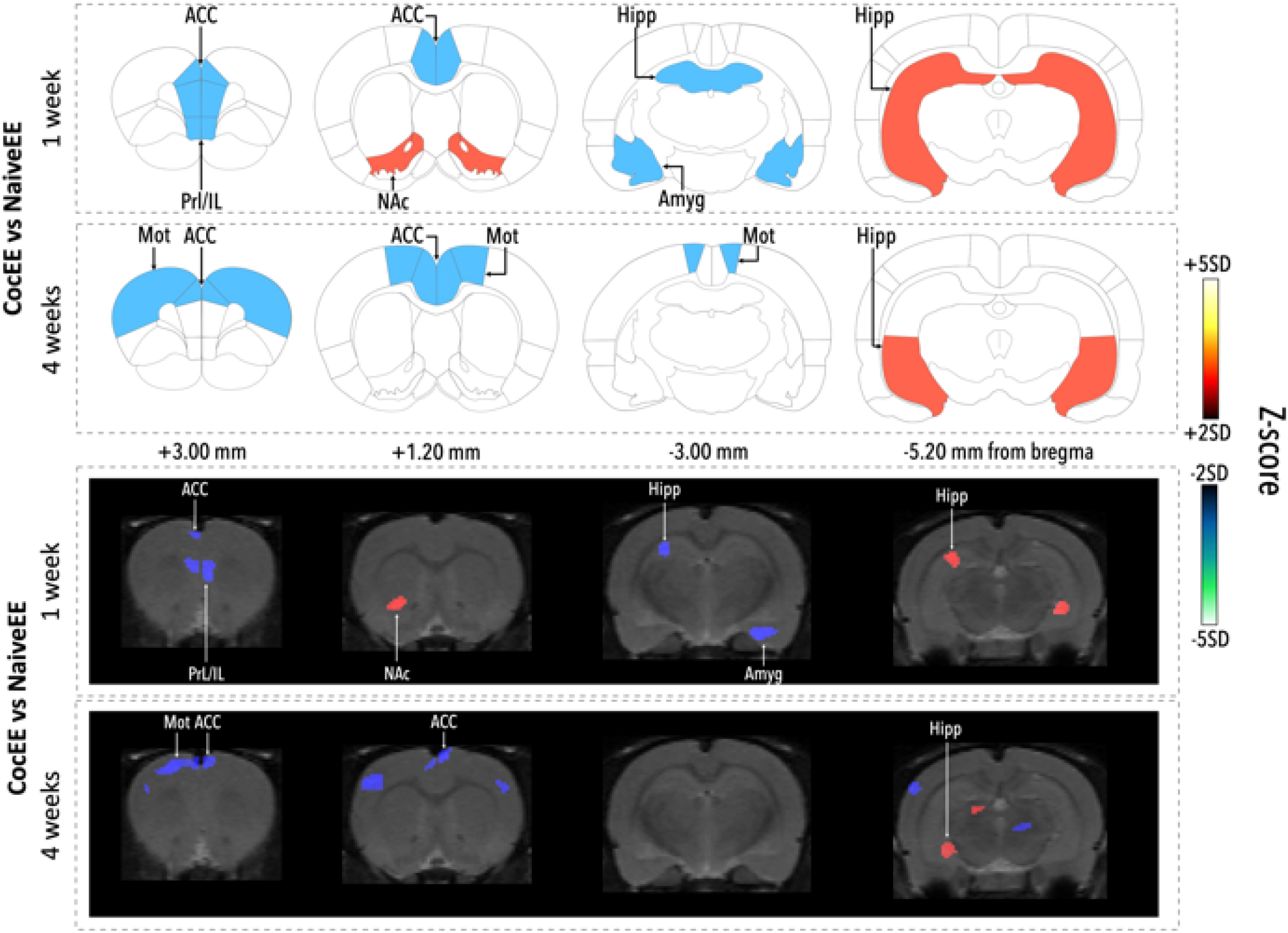
Effects of cocaine self-administration on brain metabolic activity after one and four weeks of housing in EE. Changes in metabolic activity in rats that self-administered cocaine (CocEE) compared to naïve rats (NaiveEE) and were housed in enriched environments during abstinence presented on representative coronal plates of the Paxinos and Watson atlas (upper panel), and on coronal images of z-score maps fused with an MRI template (lower panel). Increases in 18FDG uptake from dark red to yellow, decreases in 18FDG uptake from black to light blue; Student’s two-tailed t-test; p < 0.01. ACC, anterior cingulate cortex; Amyg, amygdala; Hipp, hippocampus; Mot, motor cortex; NAc, nucleus accumbens; PrL/IL, prelimbic/infralimbic cortices.

Direct comparison between cocaine-experienced rats in EE versus SE (CocEE vs CocSE, table 4 and Figure 5) revealed that after one week, EE rats showed higher metabolic activity in the ACC (p=0.0042), OFC (p=0.0003), insula (p=0.0017), dorsolateral striatum (p=0.0092), and NAc (p=0.0005), with lower activity in the amygdala (p=0.0052), anterior hippocampus (p=0.0288), and SN/VTA (p=0.0485). After four weeks, differences persisted in the OFC (p=0.0028) and dorsolateral striatum (p=0.0039), while the ACC showed reversed activity (p=0.0257) and new differences emerged in the ventral posterior hippocampus (p=0.0379).

**Table 4.** Statistically significant differences in 18FDG uptake between cocaine self-administered rats housed in EE vs SE for one and four weeks of abstinence (CocEE vs CocSE).

|  | 1 week |  |  | 4 weeks |  |  |
| --- | --- | --- | --- | --- | --- | --- |
|  | *Z*-score | *d*-value | *p*-value | *Z*-score | *d*-value | *p*-value |
| ACC | 2.84±0.57 | 1.13 | p=0.0042 | -2.24±0.82 | 0.97 | p=0.0257 |
| Mot | — | — | — | — | — | — |
| OFC | 2.87±0.81 | 1.13 | p=0.0003 | 2.70±0.42 | 1.13 | p=0.0028 |
| PrL/IL | — | — | — | — | — | — |
| Insula | 2.75±0.65 | 1.10 | p=0.0017 | — | — | — |
| DMStr | — | — | — | — | — | — |
| DLStr | 2.61±0.56 | 1.06 | p=0.0092 | 2.60±0.44 | 1.10 | p=0.0039 |
| NAc | 2.78±0.60 | 1.11 | p=0.0005 | — | — | — |
| Amyg | -3.05±0.85 | 1.15 | p=0.0052 | — | — | — |
| Ant. Hipp | -2.53±0.53 | 1.05 | p=0.0288 | — | — | — |
| Dors. Post. Hipp | — | — | — | — | — | — |
| Vent. Post. Hipp | — | — | — | 2.35±0.15 | 1.03 | p=0.0379 |
| SN/VTA | -2.36±0.19 | 0.99 | p=0.0485 | — | — | — |

ACC, anterior cingulate cortex; Amyg, amygdala; Ant. Hipp, anterior hippocampus; DLStr, dorsolateral striatum; DMStr, dorsomedial striatum; Dors. Post. Hipp, dorsal posterior hippocampus; Mot, motor cortex; NAc, nucleus accumbens; OFC, orbitofrontal cortex; PrL/IL, prelimbic/infralimbic cortices; SN/VTA, substantia nigra/ventral tegmental area; Vent. Post. Hipp, ventral posterior hippocampus.

**Figure 5.**
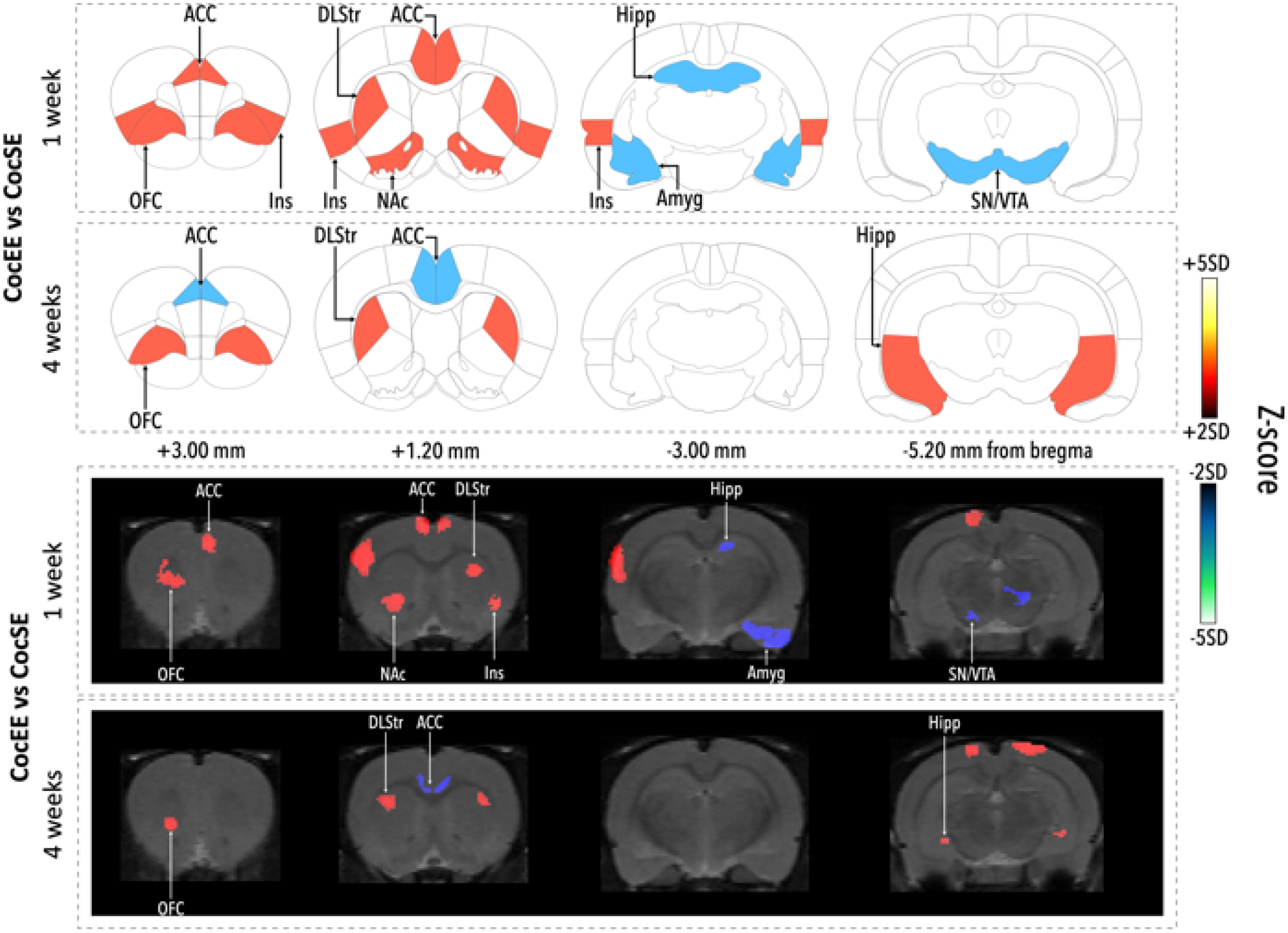
Effects of housing in EE and SE for one and four weeks on cocaine-induced metabolic changes. Changes in metabolic activity in rats that self-administered cocaine and were housed either in EE (CocEE) or SE (CocSE) during abstinence presented on representative coronal plates of the Paxinos and Watson atlas (upper panel), and on coronal images of z-score maps fused with an MRI template (lower panel). Increases in 18FDG uptake from dark red to yellow, decreases in 18FDG uptake from black to light blue; Student’s two-tailed t-test; p < 0.01. ACC, anterior cingulate cortex; Amyg, Amygdala; DLStr, dorsolateral striatum; Hipp, hippocampus; Ins, insula; NAc, nucleus accumbens; OFC, orbitofrontal cortex; SN/VTA: substantia nigra/ventral tegmental area.

## Discussion

Our findings reveal that cocaine self-administration produces persistent changes in brain metabolic activity that environmental enrichment can partially reverse. Specifically, cocaine decreased activity in cortical regions involved in executive function while increasing activity in emotional and motivational circuits. Environmental enrichment normalized activity in regions crucial for decision-making and reward processing, though some cocaine-induced changes remained unaffected.

Cocaine addiction has been shown to be associated with long-lasting neuroadaptations in brain structures and functions that are believed to play a role in the persistent risks of relapse (5,44). These changes evolve throughout the course of abstinence: some emerge during the early stages and disappear after prolonged abstinence, others persist and some only manifest after several weeks of abstinence (19,45–47). Importantly, whereas the temporary changes may reflect tolerance/withdrawal processes, those persisting after long periods of abstinence could contribute to the phenomenon of incubation of craving (48,49) and may be more relevant to relapse after treatment and detoxification (19,45–47). Consistent with our previous work (24), the changes we observed in cocaine-exposed rats closely parallel those reported in human addiction studies and align with the I-RISA model (6,7). During early abstinence, we found decreased metabolic activity in the OFC, ACC, and insula, accompanied by disrupted activity in the NAc and SN/VTA. Many of these alterations persisted through prolonged abstinence, while new changes emerged in the dorsolateral striatum and hippocampus, consistent with the development of habits and enhanced emotional processing associated with addiction (4,5,50).

In this study, environmental enrichment produced temporally distinct patterns of brain activity in naive versus cocaine-exposed animals. In naive rats, EE initially triggered widespread adaptations including decreased activity in the ACC, motor cortex, and insula, coupled with increased activity in striatal regions and hippocampus. Most of these changes normalized by four weeks, suggesting they represent acute adaptations to environmental stimulation rather than permanent circuit reorganization. These results are generally in agreement with previous studies using brain imaging approaches to investigate changes in brain activity following exposure to EE in mice (51–54) and in rats (55) and with the known molecular neuroadaptations induced by EE (56,57). In cocaine-exposed animals, however, EE demonstrated markedly different effects. Rather than producing widespread changes, EE selectively normalized cocaine-induced disruptions in specific circuits. Interesting effects were seen in the OFC and NAc, where cocaine-induced decreases in metabolic activity were prevented when animals were housed in enriched conditions. These regions are crucial for value-based decision making and reward processing (58,59), suggesting EE may restore the ability to properly evaluate natural rewards and make adaptive choices. The normalization of SN/VTA activity further supports this interpretation, as this region is critical for reward-related learning and motivation (60). Importantly, these therapeutic-like effects showed circuit specificity. While executive and reward circuits showed substantial recovery, other cocaine-induced alterations persisted despite enrichment. Most notably, the increased metabolic activity in the hippocampus remained largely unchanged, suggesting that EE may not affect all addiction-related neural adaptations equally. This selective pattern of effects provides insight into which aspects of addiction may be most amenable to environmental interventions and which may require additional therapeutic approaches.

The temporal dynamics of environmental enrichment’s effects reveal distinct phases of therapeutic action. In early abstinence, EE rapidly normalized activity in reward-related circuits, particularly the NAc and in regions involved in interoception such as the insular cortex. This rapid normalization likely reflects immediate effects on reward processing and stress response systems. The quick restoration of NAc function could explain the rapid effects of EE on drug seeking (36), possibly by enhancing the salience of natural rewards provided by enriched environment. In contrast, whereas changes in cortical regions like the OFC and in the and dorsolateral striatum were sustained from early to late abstinence, changes in the ACC evolved from increased metabolic activity at the beginning of abstinence to a decrease after four weeks of abstinence. This delayed response suggests involvement of slower neuroadaptive processes, including structural plasticity through dendritic remodeling and synaptogenesis, alterations in gene expression patterns, circuit-level reorganization, and possible restoration of normal neurotransmitter function (29,56,57). This temporal pattern suggests that successful treatment requires initial targeting of reward and stress systems while maintaining enrichment long enough for executive function recovery, accounting for different temporal windows of plasticity across circuits.

Whereas most research has investigated the neurobiological consequences of drug taking, considerably less research has been dedicated to the investigation of the mechanisms associated with recovery from addiction. Our study addresses this knowledge gap by examining the neurobiological mechanisms underlying one specific pathway to recovery—environmental enrichment. Brain imaging studies in humans and rats have demonstrated that long-term abstinence from cocaine is associated with heightened prefrontal cortical activity compared to early abstinence periods (19), which corresponds with our findings of progressive normalization of prefrontal function in EE animals. Furthermore, as discussed by Engeln & Ahmed (61), recovery can involve both the reversal of drug-induced changes and the introduction of new compensatory mechanisms that compete with drug-related circuits. Our temporal analyses align with these observations, demonstrating that EE not only reverses some drug-induced adaptations but potentially creates additional circuit-level changes that support recovery. The neuroadaptive changes we observed during EE-induced recovery likely represent just one pathway among many that can lead to remission. While our study shows that EE can reverse drug-induced neuroadaptations, particularly in the nucleus accumbens and prefrontal regions, it is important to recognize that other interventions targeting different neural mechanisms may achieve similar outcomes.

Several aspects should be considered when interpreting our results. First, our measures reflect metabolic activity under basal conditions, and therefore we cannot determine if and how environmental enrichment affects the reactivity of different brain regions to specific stimuli such as drug-related cues or stressors. Future studies using different behavioral challenges before imaging could provide insights into how environmental enrichment modifies brain responses to addiction-relevant triggers. Second, while PET imaging provides a valuable whole-brain perspective, its spatial resolution (approximately 1mm) limits our ability to distinguish small adjacent structures or subregions. For example, we analyzed the SN/VTA as a single region despite their distinct roles and could not differentiate between different amygdala nuclei or striatal subregions. More precise techniques like high-resolution fMRI or region-specific molecular analyses could complement our findings by providing finer anatomical detail. Third, to minimize the number of animals used, we performed repeated scans in the same rats. While this longitudinal approach has advantages for tracking changes over time, we cannot completely rule out potential effects of repeated manipulations such as fasting and anesthesia on brain activity. However, analysis of control rats at different time points showed no significant effects of repeated testing. Fourth, we focused only on male rats in this study. Significant differences exist in self-administration of drugs in male and female rats (62,63). On the other hand, the beneficial effects of EE on drug-related effects appears to be similar between males and females (64,65). Future studies will be needed to determine whether the neurobiological mechanisms underlying the anti-craving effects of EE are similar in male and female rats. Finally, while environmental enrichment in laboratory settings is well-defined, translation to human conditions is complex. Our enriched environment combines social, physical, and cognitive stimulation in ways that may not directly parallel human conventional environmental interventions. However, an accumulating body of literature suggests that specific aspects of enrichment such as physical activity (66,67), cognitive training (68,69) and social support (70) can be effective in the treatment of addiction. Importantly, a clinical trial is in progress to investigate the effects of combination of several of these stimulations on relapse to severe alcohol use disorder (71).

In conclusion, this study reveals that environmental enrichment can normalize cocaine-induced brain metabolic changes through distinct temporal mechanisms. Our findings provide a neurobiological framework explaining how environmental interventions can help recovery from addiction: immediate effects may reduce craving intensity by normalizing nucleus accumbens activity, while sustained environmental stimulation gradually repairs prefrontal cortical function, potentially improving decision-making and impulse control. Our results suggest that effective addiction treatment should combine early intensive intervention targeting reward and stress systems with sustained environmental support to allow executive function recovery. This multi-phase approach mirrors clinical observations that successful recovery requires both acute crisis management and long-term lifestyle changes. By understanding these neuroadaptive processes, we can design more targeted interventions that address the circuit-specific and temporally distinct aspects of addiction recovery, ultimately improving treatment outcomes for those struggling with substance use disorders.

## Acknowledgments

This work was supported by the Centre National pour la Recherche Scientifique, the Institut National de la Santé et de la Recherche Médicale, the University of Poitiers, the Nouvelle Aquitaine CPER 2015-2020 / FEDER 2014-2020 program “Habisan” and the Fondation pour la Recherche Medicale (FRM, DPA20140629806 grant, PI: M. Solinas).

We thank Miriam Melis and Celine Nicolas for helpful comments on a previous version of this manuscript.

This study has benefited from the facilities and expertise of PREBIOS platform (Université de Poitiers).

